# A global indicator of utilised wildlife populations: regional trends and the impact of management

**DOI:** 10.1101/2020.11.02.365031

**Authors:** Louise McRae, Robin Freeman, Jonas Geldmann, Grace B. Moss, Louise Kjær-Hansen, Neil D. Burgess

## Abstract

The sustainable use of wildlife is a core aspiration of biodiversity conservation but is the subject of intense debate in the scientific literature as to how, and whether, species are best used and managed. While both positive and negative outcomes of sustainable use are known for specific taxa or local case studies, a global and regional picture of trends in wildlife populations in use is lacking. We use a global data set of over 11,000 time-series to derive indices of ‘utilised’ and ‘not utilised’ wildlife populations and assess global and regional changes, principally for mammals, birds and fishes. We also assess whether ‘management’ makes a measurable difference to wildlife population trends, especially for the utilised species populations. Our results show that wildlife population trends globally are negative, but with utilised populations tending to decline more rapidly, especially in Africa and the Americas. Crucially, where utilised populations are managed, using a variety of mechanisms, there is a positive impact on the trend. It is therefore true that use of species can both be a driver of negative population trends, or a driver of species recovery, with numerous species and population specific case examples making up these broader trends. This work is relevant to the evidence base for the IPBES Sustainable Use Assessment, and to the development of indicators of sustainable use of species under the post-2020 Global Biodiversity Framework being developed under the Convention on Biological Diversity.

## Introduction

The direct use of wild species is one of the ways in which biodiversity is fundamental to the livelihoods of people (Hutton and Leader-Williams 2003; Díaz, Demissew et al. 2015; IPBES 2019). Consequently, any unsustainable impact of anthropogenic activity on species, particularly those that are important for people’s livelihoods or wellbeing, presents a threat not just to conservation but to human health and development (Pascual, Balvanera et al. 2017). The importance of the sustainable use of resources has been recognised as central to biodiversity conservation and is embedded in international bodies and conventions for nature (United Nations 1992; Hickey 1998; IUCN 2000; United Nations General Assembly 2015; IPBES 2018). However, progress towards achieving the sustainable use of resources globally remains a challenge. Progress towards Aichi target 4.2 on use within safe ecological limits was assessed as ‘poor’ in the final decadal review of the success of the strategy plan for biodiversity 2010-2020 (Secretariat of the Convention on Biological Diversity 2020) and impacts of hunting are thought to be increasing (Gallego-Zamorano, Benítez-López et al. 2020). Whilst land use change is the predominant driver of terrestrial biodiversity decline and is expected to increase in many areas (Kehoe, Romero-Muñoz et al. 2017; IPBES 2019), overexploitation is also a highly prevalent threat (Joppa, O’Connor et al. 2016) with evidence showing that harvesting, logging, fishing and hunting often occur at unsustainable levels (IPBES 2019). Together with land use change, hunting has had negative impacts on species populations, particularly in the tropics where anthropogenic pressure is currently highest and intensifying (Venter, Sanderson et al. 2016). These combined pressures have reduced the distribution of terrestrial tropical mammals, with large-bodied species the most impacted (Gallego-Zamorano, Benítez-López et al. 2020). The effect of hunting, especially for commercial use, has been implicated in causing overall decline in population abundance of 97 tropical bird and 254 tropical mammal species (Benítez-López, Alkemade et al. 2017), and the global assessments of 301 terrestrial mammals threatened with extinction list hunting as a primary threat (Ripple, Abernethy et al. 2016). In the marine realm, the percentage of commercial fish stocks that are within biologically sustainable levels decreased from 90% to 65.8% between 1974 and 2017 (FAO 2020), although recent trends suggest that stocks which are scientifically assessed are now increasing on average and intensively managed stocks are faring better (Hilborn, Amoroso et al. 2020).

The role of wildlife management is also evident in some notable examples on land. The rise of Community-Based Natural Resource Management over 30 years ago, which may include managing the use of species in place of more centralised wildlife management policies, has yielded examples of both economic and ecological benefits in many countries worldwide (Roe, Nelson et al. 2009; Anderson and Mehta 2013; Cooney, Roe et al. 2018). In regions where utilised species, particularly mammals, have been heavily impacted over centuries (Ceballos and Ehrlich 2002; Laliberte and Ripple 2004), conservation action has been implemented to stem unsustainable use and promote recovery of populations. Arguably there have been some successes with recoveries in many bird and mammal species in Europe from legal protection and habitat restoration (Deinet, Ieronymidou et al. 2013). In North America, examples of conservation and wildlife management efforts mean that once depleted populations have recovered to a level where they can be sustainably used e.g. North American bison (Sanderson, Redford et al. 2008).

As the examples above suggest, there is both evidence of successful instances of species use and of negative impacts. We propose that global and large regional views are now needed to understand how species in use are faring at scale, to measure progress towards policy targets and for identifying trends in resources that are important for people. Developing a biodiversity indicator based on species in use could fulfil these aims and also inform global processes such as the IPBES thematic assessment of sustainable use of wild species (IPBES 2018) and the development of indicators for the post-2020 global biodiversity framework. To date, synthesis of trends in species in use (herein ‘utilised species’ or ‘utilised populations’) has largely been done at the species level e.g. (Butchart 2008; Tierney, Almond et al. 2014), which may have overlooked spatially heterogeneity of impacts of use, as has been identified for commercial harvesting (Di Minin, Brooks et al. 2019). A population-based approach with information on utilisation at the site-level could provide insight that is not available at the level of species assessments and would allow small scale information on use, threats and management to be incorporated into the analysis.

To follow this approach, we develop an indicator of utilised vertebrate populations following the method used to calculate the Living Planet Index (LPI) (Loh, Green et al. 2005; Collen, Loh et al. 2009; McRae, Deinet et al. 2017), a multi-species indicator based on population trends of vertebrates used to monitor progress towards international and national biodiversity targets (Butchart, Walpole et al. 2010; Tittensor, Walpole et al. 2014; Green, McRae et al. 2020). We explore differences in these trends with respect to taxonomic groups and IPBES regions and test the sensitivity of the indicator to data quality. The Living Planet Database that sits behind the index can be disaggregated geographically and thematically at the population-level, which enables within-species comparisons and the identification of correlates predicting trends e.g. (Collen, McRae et al. 2011; Hardesty-Moore, Deinet et al. 2018). This is the basis for the second part of our analysis to contrast trends in utilised populations with those that are not used, for the complete set of species in the data set and for only those species with data for both utilised and non-utilised populations (“matched”). Finally, we explore the role that targeted management has in predicting populations trends in utilised populations.

## Results

### Geographic, taxonomic and threat data summary

The final data set comprised 11,123 population time-series from 2,944 species, of which 5,811 populations from 1,348 species were coded as utilised and 5,312 populations from 1,996 species were coded as not utilised (Table S1). In terms of utilised populations, most data were available for fish (n = 3,233) followed by mammals (n = 2,098), birds (n = 331), reptiles (n = 142) and amphibians (n = 7). Fish and mammals had more utilised populations than not, while the reverse was true for birds, reptiles and amphibians (Table S1). Geographically, our sample contained data from all IPBES regions and from 146 countries (Figure 1; see also Table S2). Both utilised and not-utilised populations were found in all regions but noticeable clusters of more utilised populations in parts of Africa, Central Asia and Canada. The largest regional data set was for the Americas. Results for Africa are based on the smallest data set of the regions; data availability throughout the time-series dropped after 2012 so the indices were shorter than the other regions, finishing in 2015 and 2013 for terrestrial/freshwater and marine respectively.

**Figure 1.**
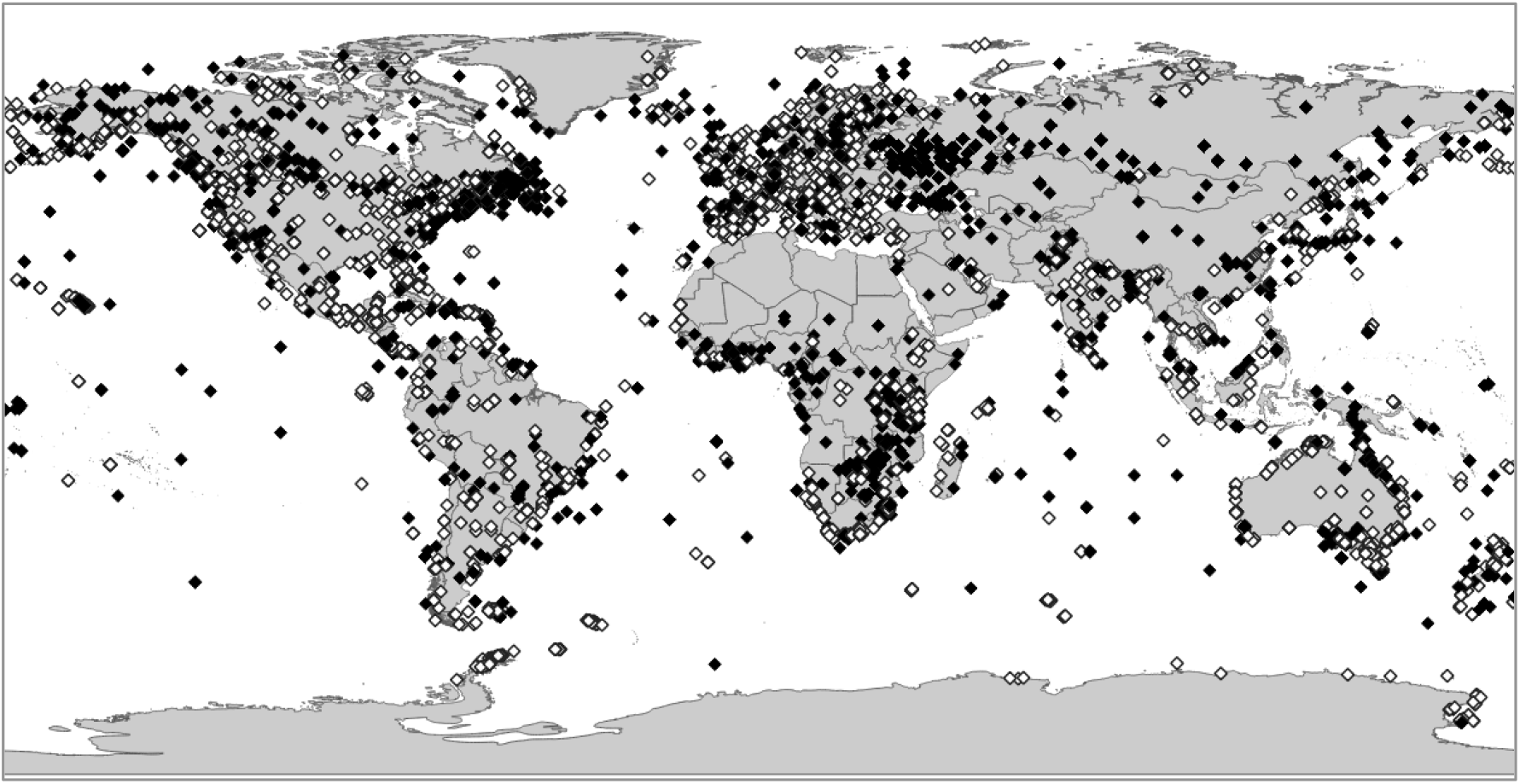
Locations of populations used in the analysis. The point location is shown for the utilised (black diamonds) and non-utilised (white diamonds) populations used in the analysis. See also Table S2

Threat information was available for 3,195 populations, 1,694 utilised and 1,501 not utilised (Table S3). There was a difference in the distribution of threats coded between utilised and not utilised populations, with a greater proportion of threats listed as Overexploitation for utilised populations (Figure S1). Nearly three-quarters of the Overexploitation threats coded for utilised populations were as a result of hunting and collecting (Figure S2). Of the utilised populations, 46% had information available on targeted management and 23% were unmanaged (remainder had no information; Table S4).

### Global indicators for utilised populations show average declines

The index for utilised populations shows a decrease of 69% for terrestrial and freshwater populations (Figure 2. Index value in 2016: 0.31; range: 0.21 to 0.44) and a decrease of 34% for marine populations (Figure 2. Index value in 2016: 0.66; range: 0.52 to 0.85), between 1970 and 2016. While the overall trend for utilised populations showed a steep decline, this was not the case for all individual populations, with 46.3% showing an overall increase, 48.9% showing an overall decrease and 4.8% were stable in the terrestrial and freshwater index. In the marine index, 53.2% of utilised populations showed an overall decline, 42.6% an overall increase and 4.2% were stable.

**Figure 2.**
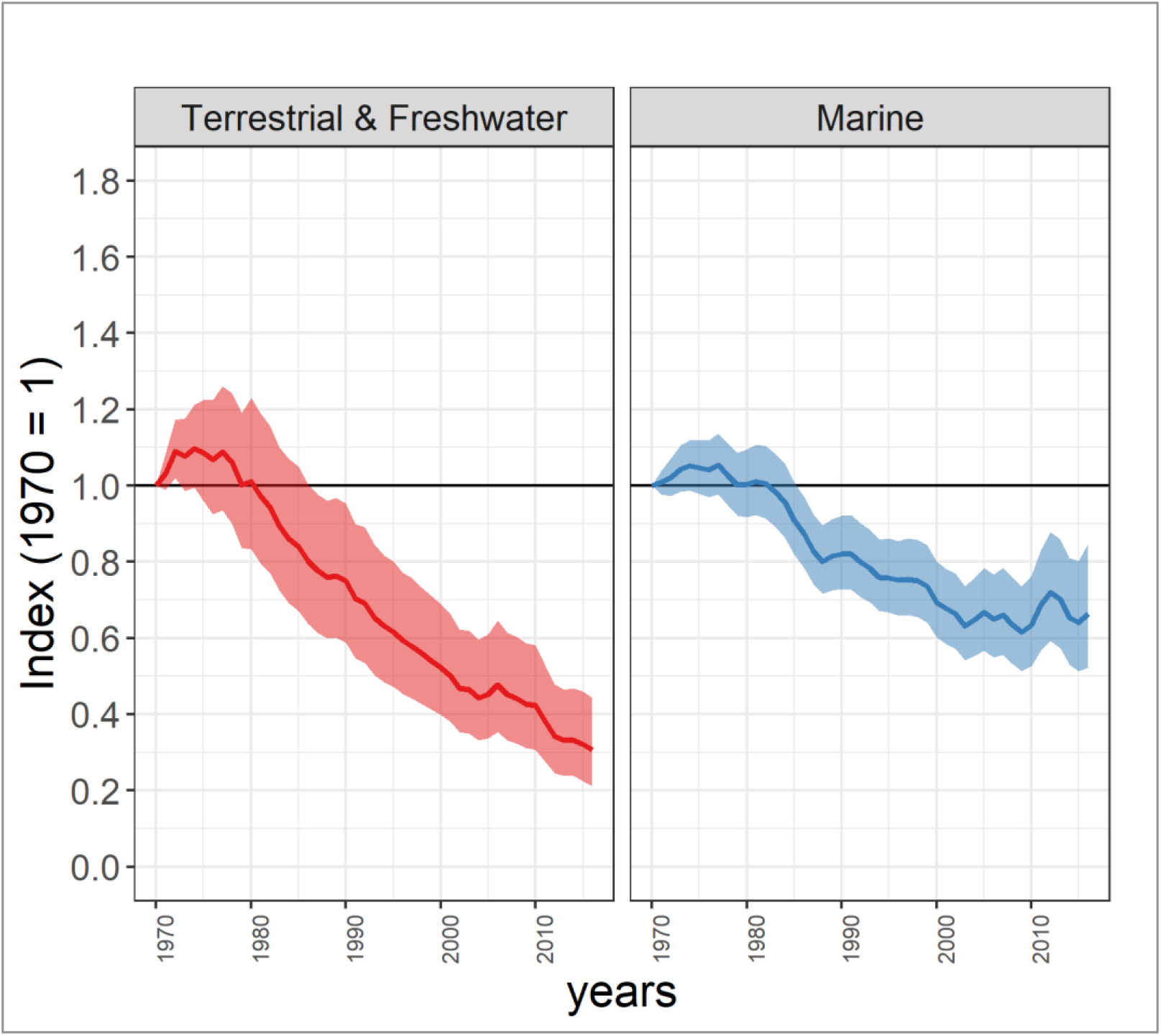
Index of utilised populations globally from 1970 to 2016. Terrestrial and freshwater index: −69%; nspp = 607, npop = 3123. Marine index: −34%; nspp = 761, npop = 2688. See Table S5 for confidence intervals.

We tested the robustness of the indices to time-series length, an important check when using population trends which vary in sample duration (Wauchope, Amano et al. 2019). We observed whether similar rates of decline were seen when restricting the data set to different thresholds for the minimum time-series length in numbers of years. When a more stringent minimum threshold for time-series length was applied, similar rates of declines were observed for the indices with a minimum of 5 years and shallower declines reported for the indices with a minimum of 10 years (Figure S3).

### Regional indices show disparate trends

The indices for utilised populations trends since 1970 by IPBES regions show disparate trends, with the largely tropical regions faring worse than the more temperate (Figure 3). The Africa indices show the greatest decline among the regions since 1970 in both the terrestrial/freshwater and marine subsets (Figure 3. Terrestrial/freshwater index value in 2015: 0.07; range 0.03 to 0.16; Marine index value in 2013: 0.08; range 0.04 to 0.17). The Asia-Pacific index shows a near-continuous decline in the marine index from 1970 to 2016 and an 83% overall decline (Figure 3. Index value in 2016: 0.17; range 0.09 to 0.31); there is a lot of uncertainty surrounding the index values in the terrestrial and freshwater index which fluctuate and ends at a similar baseline value to 1970 (Figure 3. Index value in 2016: 1.07; range 0.31 to 3.76). The terrestrial/freshwater index for the Americas showed an overall decrease of 67% between 1970 and 2016 (Figure 3. Index value in 2016: 0.33; range 0.19 to 0.58), whereas the marine index fluctuated throughout the time-series and ended at a similar baseline value to 1970, with no significant overall change (Figure 3. Index value in 2016: 1.07; range 0.78 to 1.45). The marine index for Europe and Central Asia showed a slow increase for most of the time-series after an initial decline, ending in an overall increase of 41% between 1970 and 2016 (Figure 3. Index value in 2016: 1.41; range 0.95 to 2.13). The terrestrial/freshwater index had a fluctuating trend for most of the time period but ended with a recent decline (Figure 3. Index value in 2016: 0.76; range 0.43 to 1.30).

**Figure 3.**
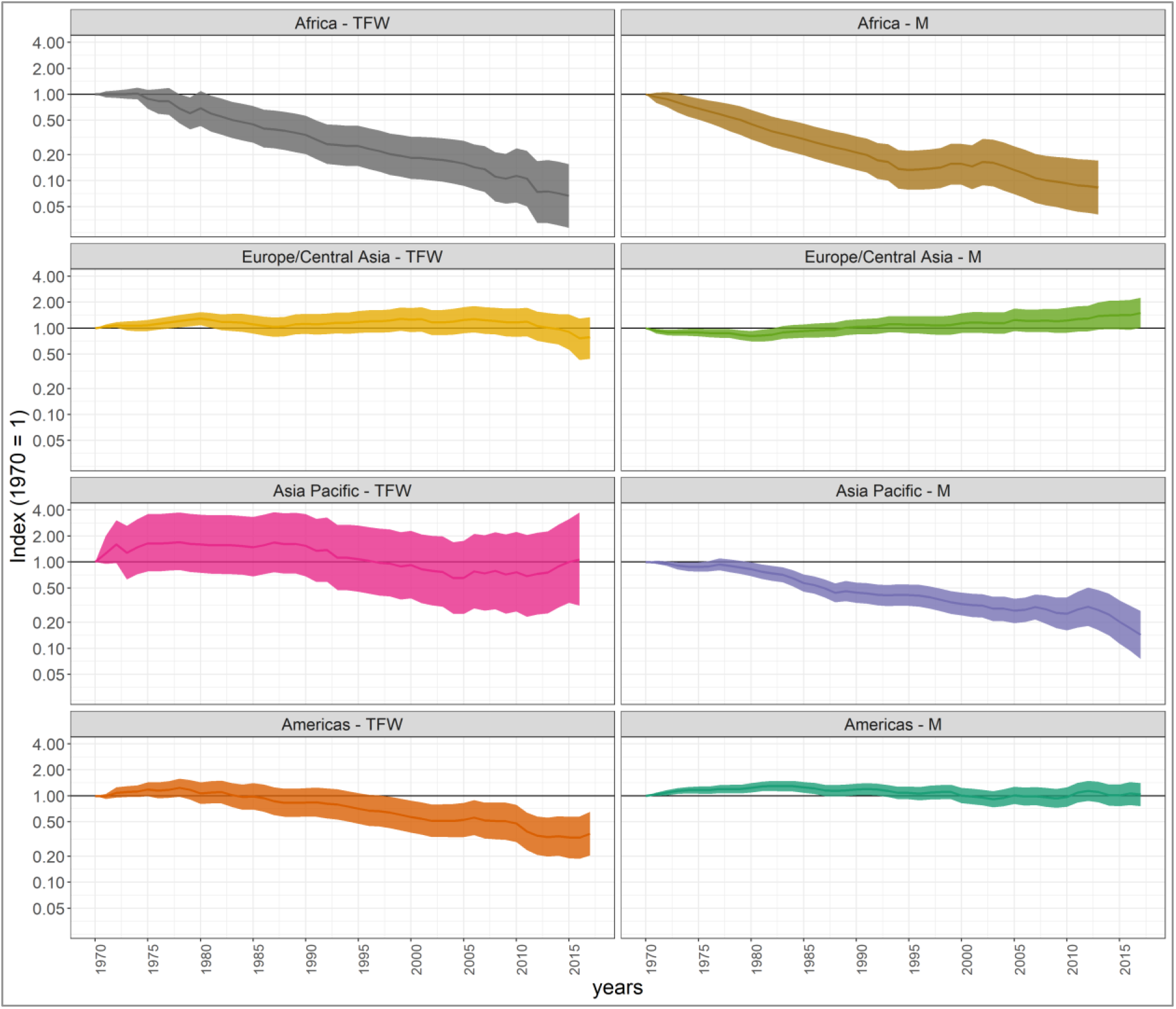
Index of utilised populations for IPBES regions from 1970 to 2016. Terrestrial and freshwater (TFW) indices: Africa (−93%; nspp = 110, npop = 314), Europe and Central Asia (−24%; nspp = 124, npop = 1886), Asia Pacific (+7%; nspp = 166, npop = 286), Americas (−67%; nspp = 239, npop = 637). Marine (M) indices: Africa (−92%; nspp = 77, npop = 132), Europe and Central Asia (+21%; nspp = 100, npop = 252), Asia Pacific (−83%; nspp = 204, npop = 349), Americas (+7%; nspp = 465, npop = 1852). See Table S5 for confidence intervals.

The subregional results for utilised populations showed either less negative or more positive trends compared to the regional index in the case of Southern Africa and North America, both terrestrial/freshwater and marine, and Central and Western Europe terrestrial/freshwater (Figure S4).

### Utilised populations show more negative trends than non-utilised on average

To explore the effect of utilisation we removed all reptile and amphibian data as these two taxa contained low number of species and populations in general but particularly those that are in the utilised category, and a large proportional difference compared to those that are not. This is likely to make unbalanced comparisons especially when dividing the data set into systems (Table S1). The remaining taxa – mammals, birds and fish – show a more stable trend for populations that are not utilised, with index values above the 1970 baseline throughout the time-series, except for a recent decline, resulting in an overall decrease of 3% over the time period (Figure 4A. Index value in 2016: 0.97; range 0.80 to 1.18). In comparison, the index for utilised populations for these taxa shows a similar trend to the index including amphibians and reptiles with an overall decline of 50% (Figure 4A. Index value in 2016: 0.50; range 0.41 to 0.62). After 1985, there is no overlap in the confidence intervals of each index which means they are significantly different.

**Figure 4.**
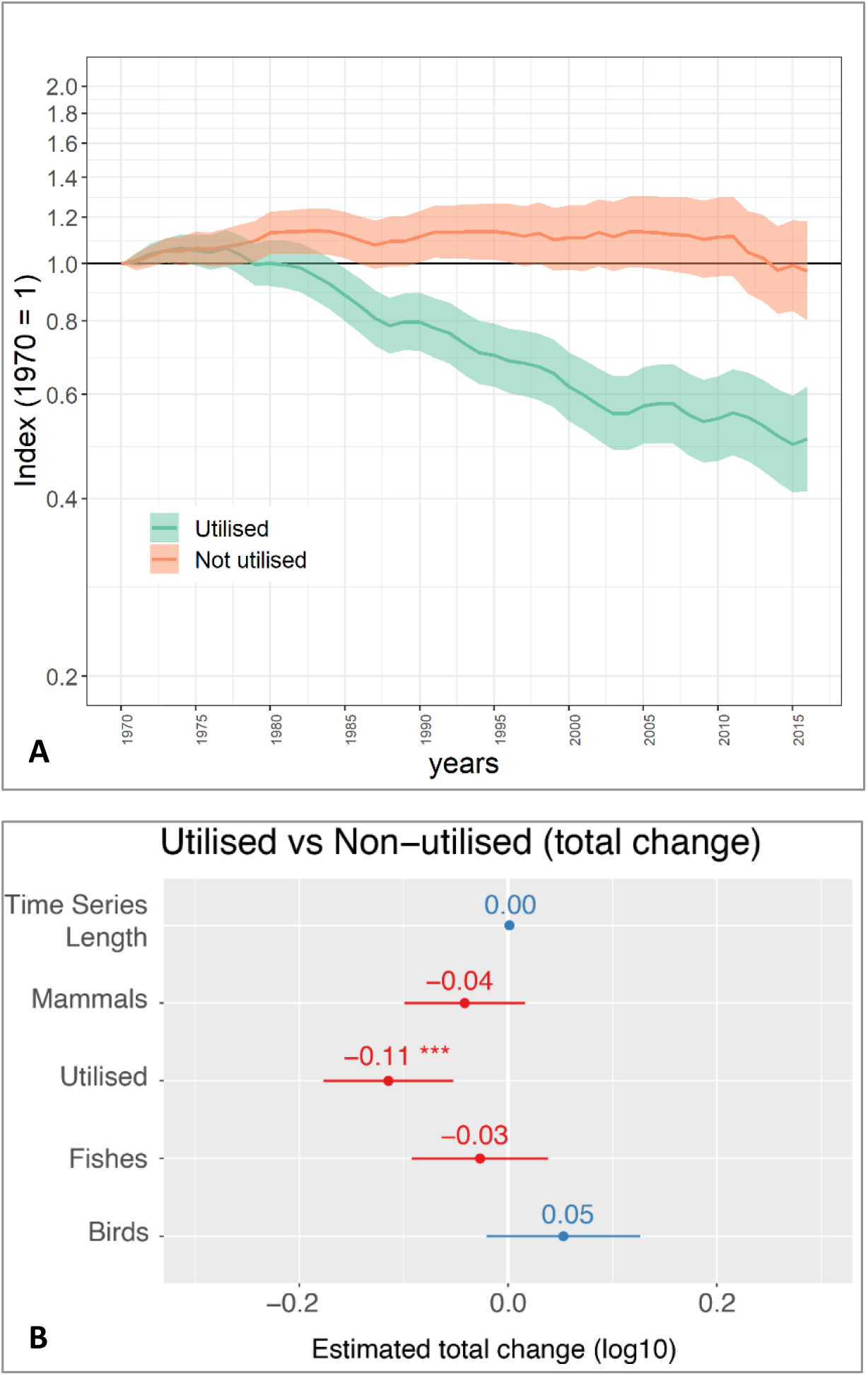
Comparison of trends in utilised and non-utilised populations from 1970 to 2016. (A) Index of utilised and non-utilised populations for species of bird, mammal and fish. Between 1970 and 2016, on average, utilised populations had declined by 50% (0.41 - 0.62) and non-utilised populations had declined by 3% (0.80 - 1.18). (B) Estimated overall total change from the best linear mixed-effect model including Family, Binomial and location as random effects. Coefficients show the estimated overall change (log10) in each group. We found no significant interaction between taxonomic group and utilisation, with utilised populations of any taxa (*Utilised*) significantly more likely to be in decline.

### Utilisation found to be a useful predictor of population trends

Utilisation was a predictor of overall population trends, with utilised populations more likely to be in decline than non-utilised. Removing utilisation from our models produced significantly worse predictions of population trends (∆AIC = −10, *χ^2^* = 11.835, p < 0.01). We found no significant interaction between utilisation and taxonomic group (∆AIC = 2, *χ^2^* = 2.0449, p = 0.3597), suggesting that all taxonomic groups are impacted by utilisation. Using our most comprehensive dataset (Mammals, Birds, Fish in Terrestrial, Freshwater and Marine systems) suggests that overall and regardless of utilisation; bird populations are slightly increasing, fish populations are generally stable, while mammals are in decline (Figure 4B). The length of a population time-series has no clear positive or negative effect on overall population trends.

We explored two taxonomic refinements to this dataset. The first removed marine fish which may represent groups of species that are under particular utilisation pressure and management. However, after removing marine fish, our results show the same pattern with utilised populations in more significant decline, (Figure S5). Our second taxonomic refinement explored these effects on terrestrial and freshwater birds and mammals (excluding all fish). Again, the results are consistent, with utilization predicting greater significant declines in population abundance (Figure S6).

As our classification of utilisation is at the population level, this may result in our models comparing groups of different species (e.g. all utilised populations may be different species to those that are not utilised). We therefore also explored a further refinement of the data only including terrestrial and freshwater bird, mammal and fish species for which we had *both* utilised and non-utilised populations (2,622 populations of 184 species. Figure S7). The comparison of trends between utilised and not utilised indices shown in Figure 4A largely holds when the trends for “matched” species are compared, although there is considerable overlap in confidence intervals until the final ten years of the time-series (Figure S8A). The mixed model result shows that utilised population trends are generally less positive than non-utilised, and there is a significant interaction between utilisation and class (Figure S8B). We also note that in this case the random effects were reduced to *Family* and *Location* (to avoid singular model fits).

### Managed populations that are utilised are less likely to be declining

For those species where we also record whether the populations are under some form of management, we find that utilised populations are less likely to be declining when management actions are in place (Figure S9). Our models suggest that - within our limited data - managed, utilised populations may be stable, but unmanaged, utilised populations are more likely to be in decline. However, we note a large taxonomic variation in these population trends, and that many populations with unknown management status have been removed.

## Discussion

### Global and regional trends in utilised populations

We present the first global indicators of trends in utilised vertebrate populations. The indices show there has been an average decline globally in monitored utilised populations between 1970 and 2016, with a starker trend amongst terrestrial and freshwater populations compared to marine. Contrasting utilised populations with those that are not utilised, we see a clear difference in the trend between the groups and this result largely holds when the same species are compared. Mixed effects models show that utilisation is a significant predictor of a more negative overall population trend. This result is robust across taxonomic subsets of our data. Whilst populations that are not utilised may be affected by threat processes such as habitat loss, it appears that the impact of utilisation in addition to the presence of other threats here is significant, as suggested in other studies (Benítez-López, Alkemade et al. 2017; Gallego-Zamorano, Benítez-López et al. 2020). Crucially, we found a positive effect of management on utilised populations, suggesting that this is a vital factor in ensuring sustainability for wildlife and people’s livelihoods.

On average, monitored populations that are utilised are in decline, according to the results presented here. This suggests that, on average, use among these populations may not be sustainable given that long-term declines are indicative of unsustainable use (Sutherland 2001). However, the global average masks some interesting variation as just under half of utilised populations had a stable or increasing trend over the time period. This implies that for some populations the use may be sustainable (according to population trend only), and that uncovering explanatory factors behind the trends is crucial. Even within a very limited suite of species for which we have both utilised and non-utilised populations and so can compare between the two, the utilised populations exhibited an overall downward trend compared to the non-utilised (Figure S8). We note, however, that this limited suite does not include species that are *only* utilised or for which we have no data on their utilisation in our dataset.

We found regional differences in trends in utilised populations. As reported in similar regional analysis of vertebrate populations (McRae, Deinet et al. 2017; WWF 2020), we found more positive trends in Europe and Central Asia, and even more so for Central and Western Europe (terrestrial and freshwater) than in other regions. However, comparisons between regions should be interpreted with care because the baseline of 1970 set for this analysis sets relatively different starting points for the state of species abundance for each region. The baseline year chosen can be important for assessing long-term trends (Collins, Böhm et al. 2020), particularly in regions where high human impact has been prevalent over centuries. In the case of North America and Western Europe, the baseline of 1970 hides historical declines in species abundance which occurred as land use was transformed after the Industrial Revolution (Ellis, Klein Goldewijk et al. 2010); post-1970 trends may therefore show fewer declines as populations stabilise but at lower numbers. This is illustrated by changes in the intactness of mammal communities drawn from estimated historical and present distributions, which suggests that mammal intactness is still high in many tropical regions but low in Europe and some areas of North America (Belote, Faurby et al. 2020), although intactness was not always directly related to the level of human modification. Another causal factor for positive trends in Central and Western Europe is the increased legal protection for hunted species, habitat conservation and agricultural land abandonment which has provided an opportunity for wildlife to rebound in many areas of Europe (Deinet, Ieronymidou et al. 2013).

Whilst the trend for North America was less negative than the wider regional trend, there was a smaller contrast in the terrestrial and freshwater index than expected given the similar context of baselines and legal protection as in Europe. Over half of the bird and mammal (55%) and freshwater fish (54%) populations in the North America index showed a declining trend. Interestingly, other analyses of trends in the United States over a similar time period showed largely positive national trends for some big game species, although declines were seen in smaller game birds (Flather, Knowles et al. 2013). One reason for the difference is likely to be the inclusion of more species in this analysis but also that freshwater fish comprised a large proportion of the utilised data set and this may be driving the average trend to an overall decline. A decrease in abundance of fish that are part of the recreational industry is thought to be occurring in Canada (Post, Sullivan et al. 2002); more broadly it has been suggested that this sector has received less cohesive management strategies than commercial fisheries (Arlinghaus, Abbott et al. 2019) and that freshwater habitats in particular can be logistically hard to manage (Post, Sullivan et al. 2002). The decline may also be attributed to changes in habitat, especially fragmentation of river systems, which is a particular threat to migratory freshwater fish (Deinet, Scott-Gatty et al. 2020).

Data availability was a limitation to assessing trends for Asia Pacific and Africa; for the latter it was mainly an issue in the later years of the time-series. However, significant declines compared to the 1970 baseline were seen in both Africa indices and in the marine Asia-Pacific index. The extensive variability in the terrestrial and freshwater data from Asia-Pacific resulted in an inconclusive trend. The subregional analysis for Southern Africa did produce significantly less negative trends for both terrestrial and freshwater as well as marine indices, indicating that approaches to sustainable management of wildlife, including incentivised use of species may have mitigated steeper declines and promoted some populations to stables or recover, although not enough to prevent an overall decline on average. With the analysis conducted at a regional and subregional scale, the results may mask the relative differences between countries. For example, positive trends in wildlife have been shown in Namibia as a result of Community-Based Natural Resource Management (Naidoo, Weaver et al. 2011), which runs counter to the overall result for Africa and Southern Africa and illustrate that management strategies that focus on sustainable use can be successful. These regional indices therefore have the advantage of providing a large-scale indicator as an overview, but the results don’t necessarily represent trends at smaller scales and can hide many local examples of ‘best-practice’. However, the data and method described here is applicable at national and regional level (McRae, Böhm et al. 2012; WWF-Canada 2020) and could be tailored to assess trends in utilised species at difference scales provided sufficient data is available.

### Results in the context of sustainable use

The Convention on Biological Diversity definition of sustainable use as: *“the use of components of biological diversity in a way and at a rate that does not lead to the long-term decline of biological diversity, thereby maintaining its potential to meet the needs and aspirations of present and future generations”* (United Nations 1992).

Our results show a long-term decline, on average, amongst utilised populations globally suggesting use, overall, is not sustainable. This aligns with broad-scale findings of the impacts of utilisation on mammals and birds (Ripple, Abernethy et al. 2016; Benítez-López, Alkemade et al. 2017) and of trends in utilised fishes (FAO 2020; Palomares, Froese et al. 2020). However, the breakdown of utilised populations showing increasing and declining trends reveals a roughly equal split, inferring that the use of half of the populations in the data set appears to not jeopardize the long-term persistence of these population. Sustainable use as a tool is harder to analyse explicitly with this data set as the implementation of this as a tool was not recorded. However, utilised populations where the use was incentivised for conservation are likely to also be categorised as ‘managed’ due to regulations or guidance to manage the use. On this assumption, the impact of management seen on utilised populations in this analysis indicates that populations intentionally utilised in this way are likely to have more positive trends.

The incorporation of management into this analysis introduces important nuance, suggesting that less negative or more positive trends are likely if sustainable management of utilised species is pursued. Management can take many forms and utilisation itself can be a tool for both conservation and human development, providing incentives for habitat and species conservation to support the provision of resources for people into the future e.g. (Lichtenstein 2009; Fukuda, Webb et al. 2011). Harvesting of Saltwater crocodile (*Crocodylus porosus*) eggs for commercial farms in the Northern territory of Australia has been an incentive for its conservation and an increase in density indices suggested a full recovery from uncontrolled hunting since 1975 (Fukuda, Webb et al. 2011). Likewise, the establishment of communal conservancies in Namibia were found to provide dual benefits to the local community from tourism and hunting, especially when these activities occurred in parallel (Naidoo, Weaver et al. 2016); recoveries in the abundance of species have also been recorded (Naidoo, Weaver et al. 2011). Given the importance of community involvement in sustainable management (Persha, Agrawal et al. 2011; Coad, Fa et al. 2019), context is key and there may not be a single approach to take for sustainably managing wildlife for conservation and livelihood outcomes (Persha, Agrawal et al. 2011; Anderson and Mehta 2013); however some over-riding considerations, such as avoiding protectionist approaches and engaging community in decision-making, have been noted (Cooney, Roe et al. 2018).

Sustainable management has arguably had more focus in the marine realm which could offer an explanation of the more positive trends seen in the marine indices for Central and Western Europe, and North America. In response to concerns about overfishing and in light of well-documented cases of fish stock collapse, such as Newfoundland cod (Hutchings and Myers 1994) and northeast Atlantic herring (Dickey-Collas, Nash et al. 2010), efforts to manage fisheries at national and international levels began to develop in the 1970s and 80s (Sissenwine, Mace et al. 2014). Although commercial stocks are often reported as in decline globally (Palomares, Froese et al. 2020), there are studies which highlight positive trends in stocks, particularly those which have been intensively managed to avoid over-fishing (Hilborn, Amoroso et al. 2020). The nature of the global fishing industry means that global management is required for many fish stock – in particular those outside national waters. However for fisheries nearer to coastal communities, management at smaller scales, specifically community co-management, is advocated as a viable and realistic long-term solution for sustainable fishing (Gutiérrez, Hilborn et al. 2011). Importantly, this form of management is also likely to be more equitable. Successful case studies of community co-management have been found from an assessment across all regions of the world, with the best outcomes determined by attributes of the community - the presence of community leaders, strong social cohesion – and of the management approach – community-based protected areas and individual or community quotas (Gutiérrez, Hilborn et al. 2011).

### Suitability as an indicator of utilised populations

The use of the LPI data and method as the basis for an indicator for utilised populations has the advantage of capitalising on available data and a method already established in research and policy (Collen, Loh et al. 2009; Tittensor, Walpole et al. 2014; McRae, Deinet et al. 2017; Secretariat of the Convention on Biological Diversity 2020). Population trend data procures advantage over species level assessments as it incorporates site specific information on utilisation and management which can vary across a species range and over time. Abundance trends also incorporate a sensitivity meaning the index can respond quickly to changes in populations (Santini, Belmaker et al. 2017). As an indicator of populations that are important for human use, this can be a useful early-warning signal that management intervention needs to be initiated or made more effective to sustain vital resources.

A primary shortcoming of this approach is with respect to the shortage of comprehensive information for all vertebrate groups and the lack of plant or invertebrate data. The data set behind the index suffers much of the same biases as found in other data sets and indicators (McRae, Deinet et al. 2017; Proença, Martin et al. 2017), with data available for well-studied taxa such as birds and mammals, or those of commercial importance such as fish. Geographic gaps in the data also remain, particularly in South America and South-east Asia, regions that are hotspots of both wildlife trade (Scheffers, Oliveira et al. 2019) and of mammals threatened by hunting (Ripple, Abernethy et al. 2016). However, it can be prudent to develop indicators in lieu of comprehensive data, providing that the gaps in data are clear and biases addressed when feasible (Jones, Collen et al. 2011; McRae, Deinet et al. 2017).

Whilst population trend is one measure of sustainability, there are other factors which are not considered here and might not be appropriate to aggregate into a global indicator. Changes in population structure as a result of selective hunting pressure can occur e.g. (Garel, Cugnasse et al. 2007), which may start to occur prior to a population decline being detected. A utilised population may show altered behaviours e.g. (Ciuti, Muhly et al. 2012) which may not necessarily correlate with population trends. Finally, this index is not able to demonstrate what is the level of sustainable use and how far beyond this limit are current levels of pressure – i.e. how much would the current use need to be reduced to reverse the declines observed. The human dimension of sustainable use, relating to the needs and benefits of people’s use of wildlife is not factored into this analysis but is a fundamental aspect of how sustainably species are used (Hutton and Leader-Williams 2003). Dividing the utilised populations into types of use could help in this regard and incorporating socioeconomic data would be challenging but an interesting consideration to develop this indicator further. This work also does not address the non-consumptive component of utilisation. Incorporating trend data for species under this type of use might introduce more positive or stable trends, on the assumption that non-consumptive use is less likely to directly cause population decline, even though the effects of uses such as tourism could be detrimental to some species e.g. (Kelly, Pickering et al. 2003; Burgin and Hardiman 2015).

## Conclusion

The alignment of conservation and human development goals is challenge, particularly when it comes to the sustainable use of resources (Hutton and Leader-Williams 2003). The results presented here suggest that whilst the global trend is negative on average for utilised populations, evidence from a substantial data set of utilised populations suggest that managing the use of wildlife has had a positive impact on species trends. This is an important finding for the conservation of species directly of benefit to people. With sustainable use a core component of both the post-2020 Global Biodiversity Framework and the Sustainable Development Goals, indicators are required to monitor progress towards the associated targets; the index presented here can address this need.

## Experimental procedures

### Resource availability

#### Lead contact

Further information and requests for resources should be directed to and will be fulfilled by the lead contact, Louise McRae.

#### Materials availability

This study did not generate new unique materials

#### Data and code availability

The data used in this paper is stored in the online database at www.livingplanetindex.org. The R package used for analysis is available here: https://github.com/Zoological-Society-of-London/rlpi

### Collection and coding of data set

Vertebrate population time series data were extracted from the Living Planet Database (WWF/ZSL 2020), a global repository of annual abundance estimates collated primarily from the scientific literature and online databases (Collen, Loh et al. 2009; McRae, Deinet et al. 2017). The annual abundance measures were collected using a consistent monitoring method in a given and consistent location. The time-series vary from 2 to 46 years in terms of length of timeframe and in number of raw annual data points. Units of abundance were population size estimates, densities or proxies of abundance, such as nests or breeding pairs (see (McRae, Deinet et al. 2017) for more details). Alongside the abundance data for each population, several ancillary data fields were extracted to use for summaries, disaggregation and modelling of the data (Table S6).

The use of species can be consumptive - hunting, fishing, harvesting - or non-consumptive – tourism, cultural experiences, catch and release fishing – and for commercial, subsistence or recreational purposes (Sustainable Use and Livelihoods Specialist Group 2020). The definition of utilised in the Living Planet Database, refers only to consumptive use but does not include non-consumptive uses. If a population is in use as a form of management, it will be tagged as both ‘utilised’ and ‘managed’. The two categories allow us to differentiate between populations that are utilised and under management with those that are utilised and unmanaged. Additionally, we consider populations that are not utilised but are managed for some other purpose e.g. provision of nest boxes for a species whose nesting habitat has been degraded.

### Index calculation

Using the R package *rlpi* (https://github.com/Zoological-Society-of-London/rlpi) and following the Generalised Additive Modelling framework in (Collen, Loh et al. 2009), we calculated global and regional indices of abundance for populations that were utilised and populations that were not. IPBES regions were chosen to divide the data sets, as these are commonly used for reporting on broad scale biodiversity trends. The indices were calculated for different subsets of the data (Table S7). The subset of species in the data set with data for both utilised and non-utilised populations are referred to as “matched” species.

The finer scale subregional analysis was conducted for three subregions – Southern Africa, Central and Western Europe and North America. Wildlife management in these subregions has arguably been more widespread so a comparison with the wider regional trends is of interest.

The baseline year set for the index was 1970 and it was run until 2016, as data availability decreases beyond this year due to the publication time lag. Each population trend carried equal weight within each species and each species trend carried equal weight within each index; we did not incorporate any additional weighting as has been done for the global LPI (McRae, Deinet et al. 2017). The confidence intervals were calculated using bootstrap resampling of 10,000 iterations to indicate variability in the underlying species trends (Collen, Loh et al. 2009).

### Mixed models

We considered how total population abundance change (*T_lambda, sum of annual rates of change*) had changed in response to utilisation (*Utilised)* for different taxonomic groups (*Class*: Mammalia, Aves, Fishes). Time-series length was included to understand if longer population trends tended to reflect more positive or negative overall change. Taxonomic and site effects were accounted for by including a random intercept for Family, Binomial (Genus + species) and population location. *T_lambda* values were taken from the *rlpi* package, which generates a matrix of annual rates of change for each population. The annual rates were summed to give a logged value of total change in abundance for each population `(T_lambda ~ 0 + TS_length + Utilised + Class + (1|Family/Binomial) + (1|Location`).

We also explored how the removal of marine populations and fish population affected this model. For a subset of these populations we also have information on whether they are subject to some form of management. We therefore assess a second model structure including Management as an additional explanatory factor (`T_lambda ~ 0 + TS_length + Management + Utilised + Class + (1|Family/Binomial) + (1|Location)`.

## Supplemental Information

Document S1. Figures S1-S9 and Tables S1-S7

## Acknowledgements

This output has been funded in whole or part by the UK Research and Innovation’s Global Challenges Research Fund, under the Trade, Development and the Environment Hub project (project number ES/S008160/1). LM and NDB were funded by UNEP-WCMC through the above grant. The authors would like to kindly thank Dilys Roe for her insightful review of the draft manuscript.

## Author contributions

Conceptualisation, All authors; Data curation, L.M. and R.F.; Formal Analysis, L.M. and R.F.; Funding acquisition, N.D.B; Writing – original draft, L.M.; Writing – review & editing, All authors

