## Supplemental information for "A global indicator of utilised wildlife populations: regional trends and the impact of management"

**Table S1 Number of species and populations in the LPD by Class, system and utilised status**

| | | Species | | Populations | |
|---|---|---|---|---|---|
| | | Utilised | Not utilised | Utilised | Not utilised |
| **Birds** | **Freshwater** | 69 | 154 | 80 | 310 |
| | **Marine** | 15 | 189 | 39 | 707 |
| | **Terrestrial** | 60 | 442 | 212 | 849 |
| | ***total*** | **144** | **785** | **331** | **1866** |
| **Mammals** | **Freshwater** | 11 | 15 | 129 | 49 |
| | **Marine** | 27 | 55 | 76 | 211 |
| | **Terrestrial** | 164 | 356 | 1893 | 802 |
| | ***total*** | **202** | **426** | **2098** | **1062** |
| **Reptiles** | **Freshwater** | 25 | 31 | 66 | 59 |
| | **Marine** | 6 | 14 | 68 | 80 |
| | **Terrestrial** | 4 | 144 | 8 | 212 |
| | ***total*** | **35** | **189** | **142** | **351** |
| **Amphibians** | **Freshwater** | 3 | 81 | 4 | 154 |
| | **Terrestrial** | 2 | 93 | 3 | 149 |
| | ***total*** | **5** | **174** | **7** | **303** |
| **Fish** | **Freshwater** | 276 | 113 | 728 | 233 |
| | **Marine** | 713 | 404 | 2505 | 1497 |
| | ***total*** | **989** | **517** | **3233** | **1730** |

**Table S2 Number of populations in each IPBES region**

| IPBES region | Utilised | Not utilised |
|---|---|---|
| Africa | 446 | 396 |
| Americas | 2489 | 2729 |
| Asia-Pacific | 635 | 949 |
| Europe-Central Asia | 2138 | 1093 |

**Table S3 Number of populations with population-level threat information**

| Population threat status | Utilised | Not utilised |
|---|---|---|
| No threats | 390 | 1065 |
| Threatened | 1694 | 1501 |
| Unknown (large data set) | 1475 | 1386 |
| Unknown (no information) | 2252 | 1360 |

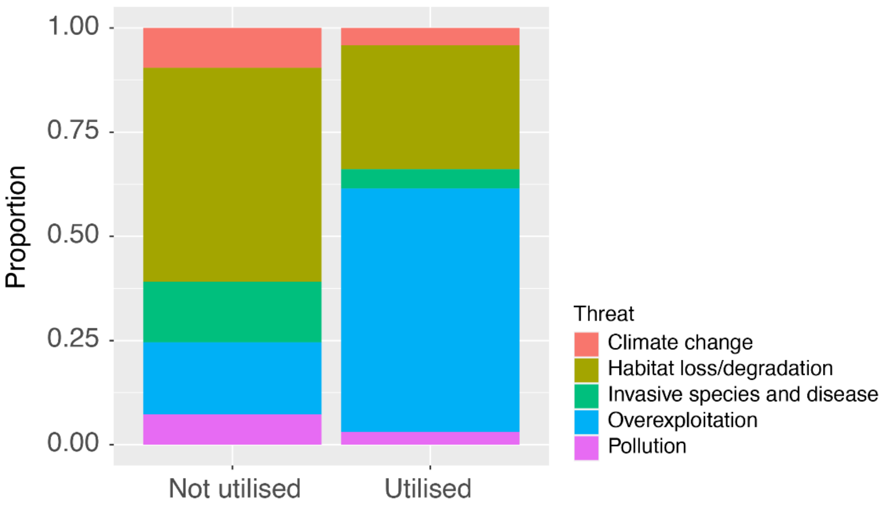


**Figure S1. Recorded threats to populations.** Populations can be not utilised but still threatened by exploitation e.g. indirect killing, persecuted as a pest

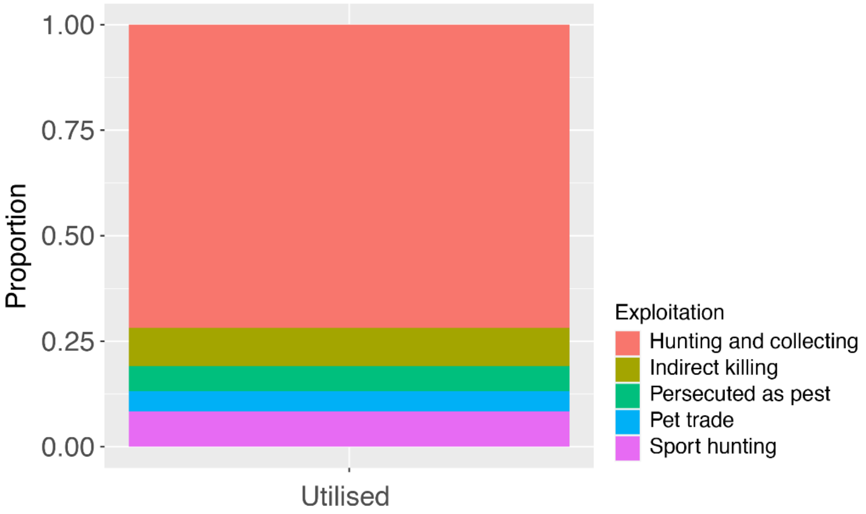


**Figure S2 Recorded categories for populations threatened by Overexploitation**

**Table S4 Number of populations with management information**

| Managed population | Utilised | Not utilised |
|---|---|---|
| **No** | 1329 | 2902 |
| **Yes** | 2671 | 704 |
| **Unknown** | 1811 | 1706 |

**Table S5 – Final Index values for Indices presented in main text, Related to Figure 2 and Figure 3.** For each trend the final index value (2016 i) is shown with the calculated lower and upper 95% bounds of that value from bootstrapping.

| Name | Final Value (2016) | Lower | Upper |
|---|---|---|---|
| Global Utilised (TFW) | 0.31 | 0.21 | 0.44 |
| Global Utilised (Marine) | 0.66 | 0.52 | 0.85 |
| Africa (TFW) | 0.07† | 0.03 | 0.16 |
| Africa (Marine) | 0.08†† | 0.04 | 0.17 |
| Europe and Central Asia (TFW) | 0.76 | 0.43 | 1.30 |
| Europe and Central Asia (Marine) | 1.41 | 0.95 | 2.13 |
| Asia Pacific (TFW) | 1.07 | 0.31 | 3.76 |
| Asia Pacific (Marine) | 0.17 | 0.09 | 0.31 |
| Americas (TFW) | 0.33 | 0.19 | 0.58 |
| Americas (Marine) | 1.07 | 0.78 | 1.45 |
| Utilised Populations (Birds, Mammals, Fish) | 0.50 | 0.41 | 0.62 |
| Non-utilised populations (Birds, Mammals, Fish) | 0.97 | 0.80 | 1.18 |
| Utilised Populations (TFW) > 5 years | 0.31 | 0.22 | 0.45 |
| Utilised Populations (Marine) > 5 years | 0.71 | 0.56 | 0.90 |
| Utilised Populations (TFW) > 10 years | 0.46 | 0.34 | 0.62 |
| Utilised Populations (Marine) > 10 years | 0.83 | 0.66 | 1.05 |
| Southern Africa (TFW) | 0.41†† | 0.17 | 0.91 |
| Southern Africa (Marine) | 0.16†† | 0.08 | 0.30 |
| C/W Europe (TFW) | 1.39 | 0.77 | 2.50 |
| C/W Europe (Marine) | 1.24 | 0.80 | 1.98 |
| North America (TFW) | 0.59 | 0.33 | 1.07 |
| North America (Marine) | 1.21 | 0.87 | 1.67 |
| Utilised Populations (Birds, Mammals, Fish; TFW) | 0.33 | 0.22 | 0.50 |
| Non-utilised populations (Birds, Mammals, Fish; TFW) | 0.81 | 0.65 | 1.01 |
| Utilised Populations (Birds & Mammals; TFW) | 0.32 | 0.20 | 0.51 |
| Non-utilised populations (Birds & Mammals; TFW) | 0.88 | 0.71 | 1.09 |
| Utilised Populations (Birds, Mammals, Fish; TFW; Matched Species) | 0.53 | 0.29 | 0.95 |
| Non-utilised populations (Birds, Mammals, Fish; TFW; Matched Species) | 2.07 | 1.18 | 3.63 |

*† 2015 value due to lack of later data; †† 2013 value due to lack of later data*

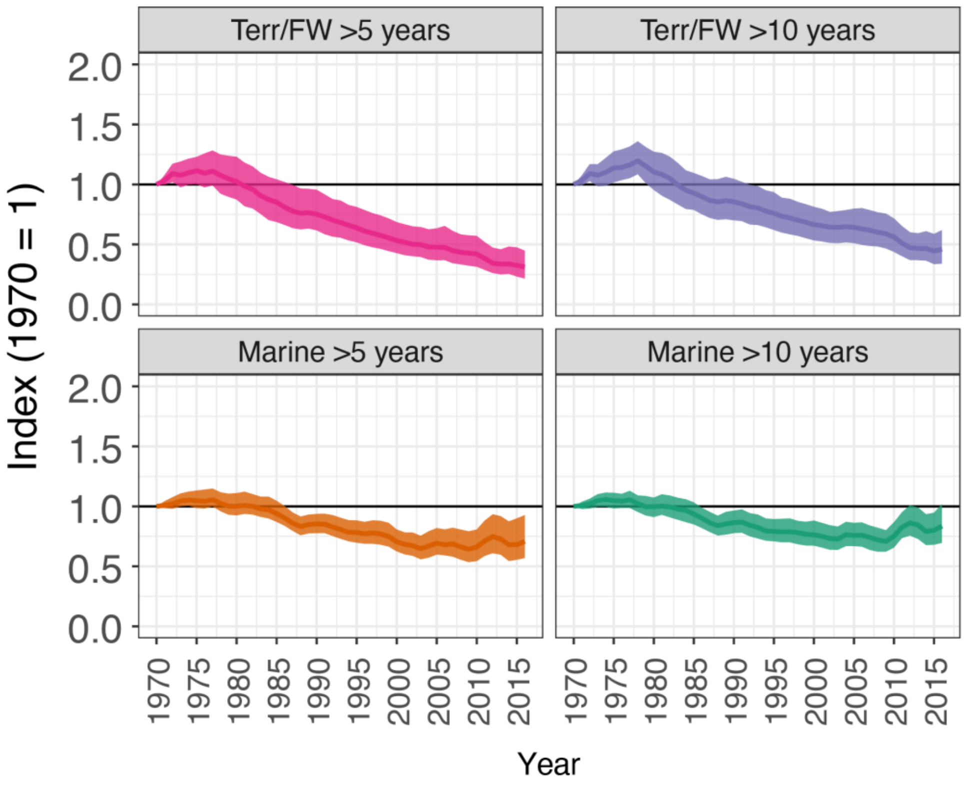


**Figure S3 Effect of time-series length on index of utilised populations, related to Figure 2.** Indices of utilised populations of terrestrial/freshwater (Terr/FW, upper) or marine (lower) for different subset of data which only include time series that span more than 5 years (>5) or more than 10 years (>10). See Table S5 for final values and confidence intervals.

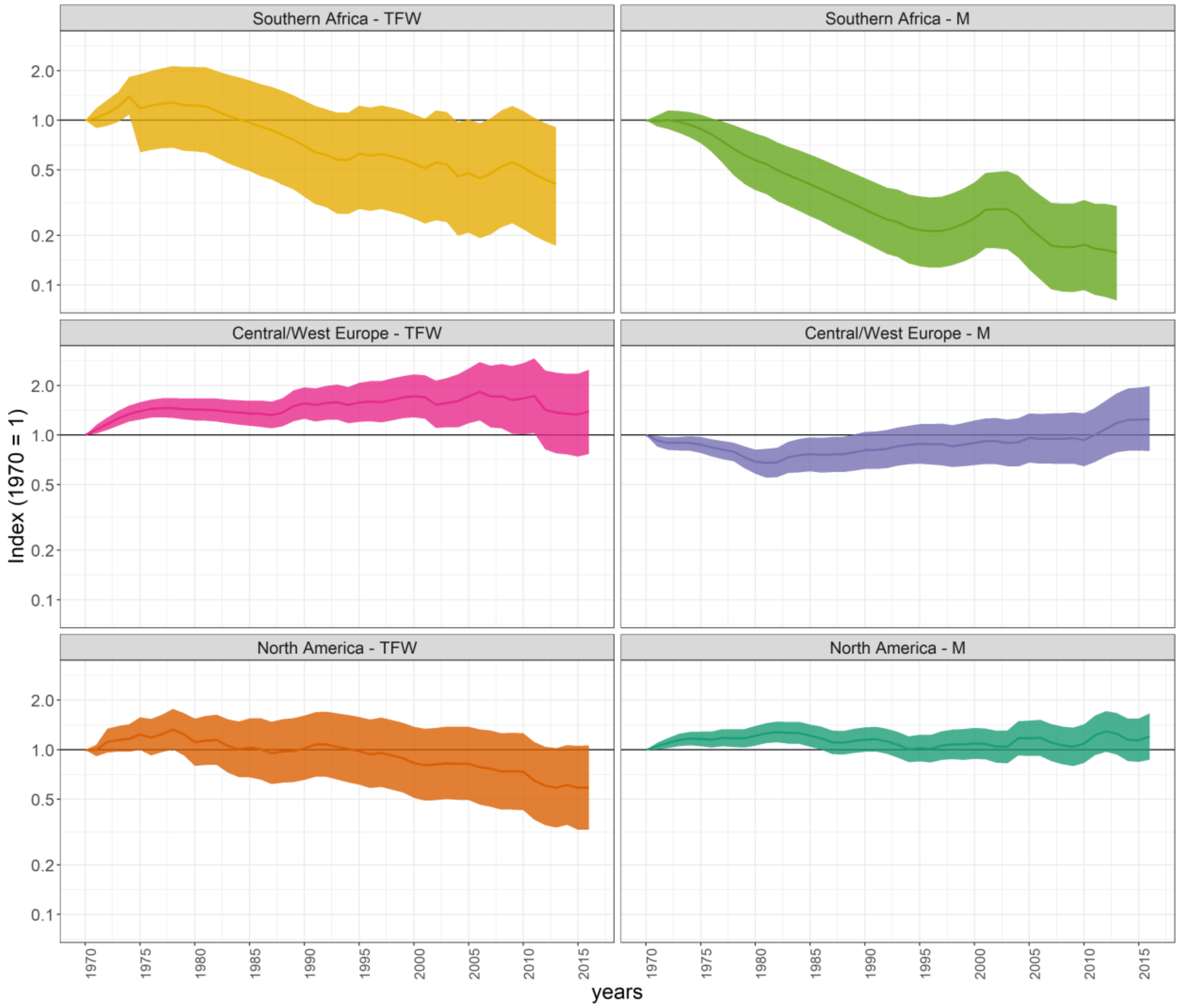


**Figure S4 Index of utilised populations for IPBES subregions, related to Figure 3.** Indices of utilised d populations of terrestrial/freshwater (TFW, left) or marine (M, right) for different IPBES subregions. See table S5 for final values and confidence intervals.

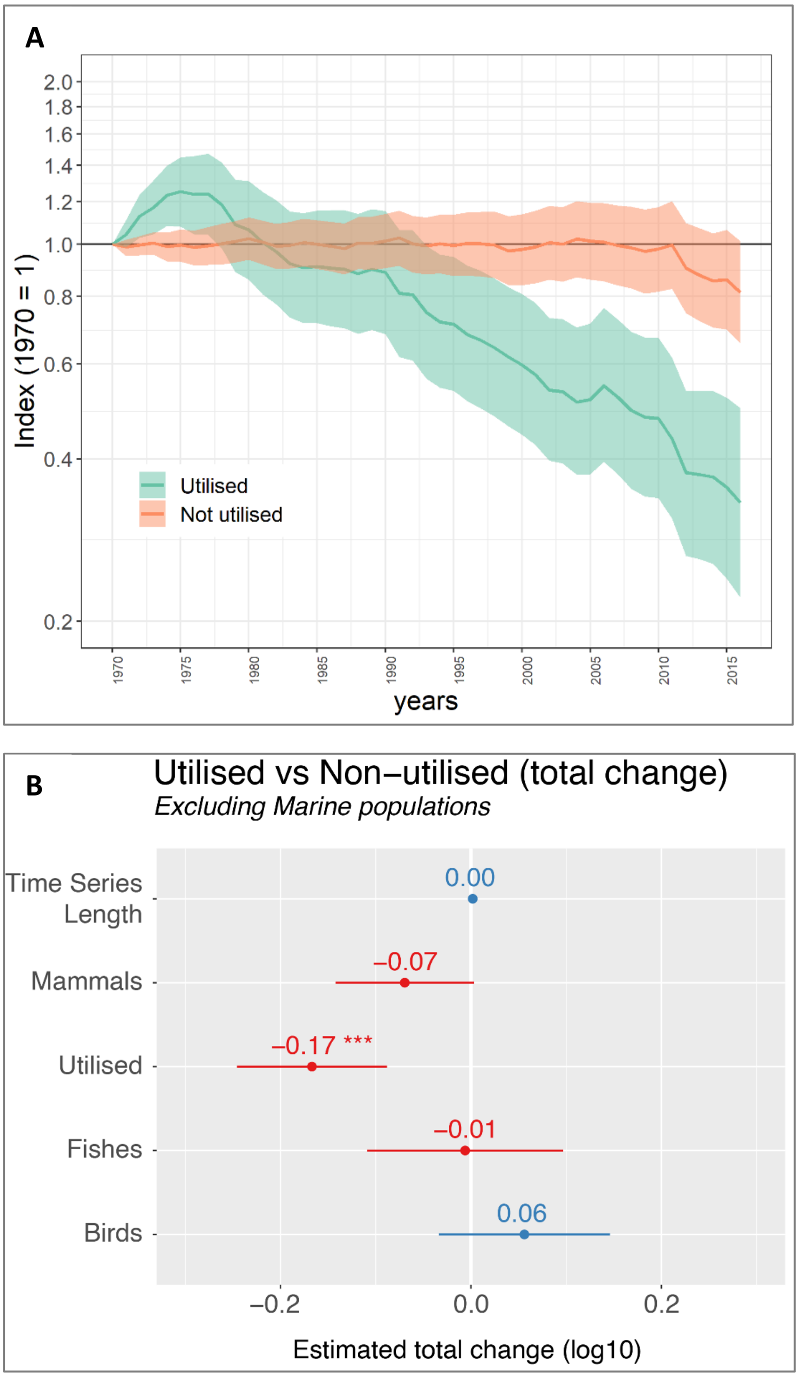


**Figure S5 Index of utilised and non-utilised populations for species of terrestrial and freshwater birds, mammals and fish; Related to Figure 4. (A)** Here, between 1970 and 2016, on average, utilised populations had declined by 67% (0.22 - 0.50) and non-utilised populations had declined by 19% (0.65 - 1.01). **(B)** Estimated overall total change from the best linear mixed-effect model including Family, Binomial and location as random effects. Coefficients show the estimated overall change (log10) in each group. We found no significant interaction between taxonomic group and utilisation, with utilised populations of any taxa (*Utilised*) significantly more significantly more likely to be in decline.

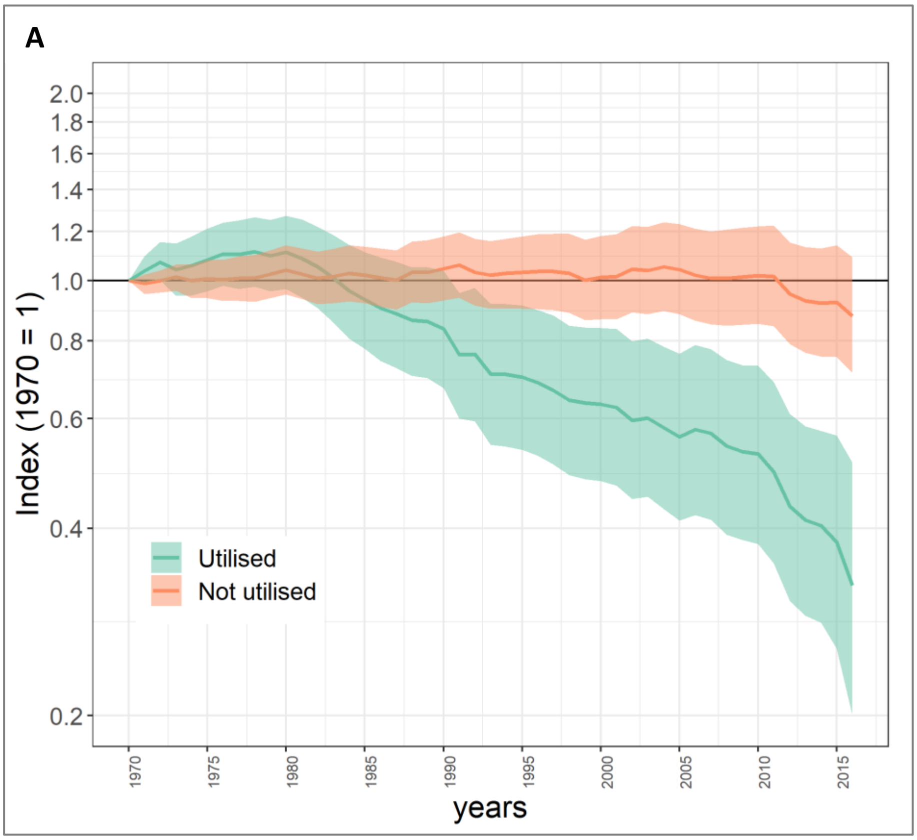


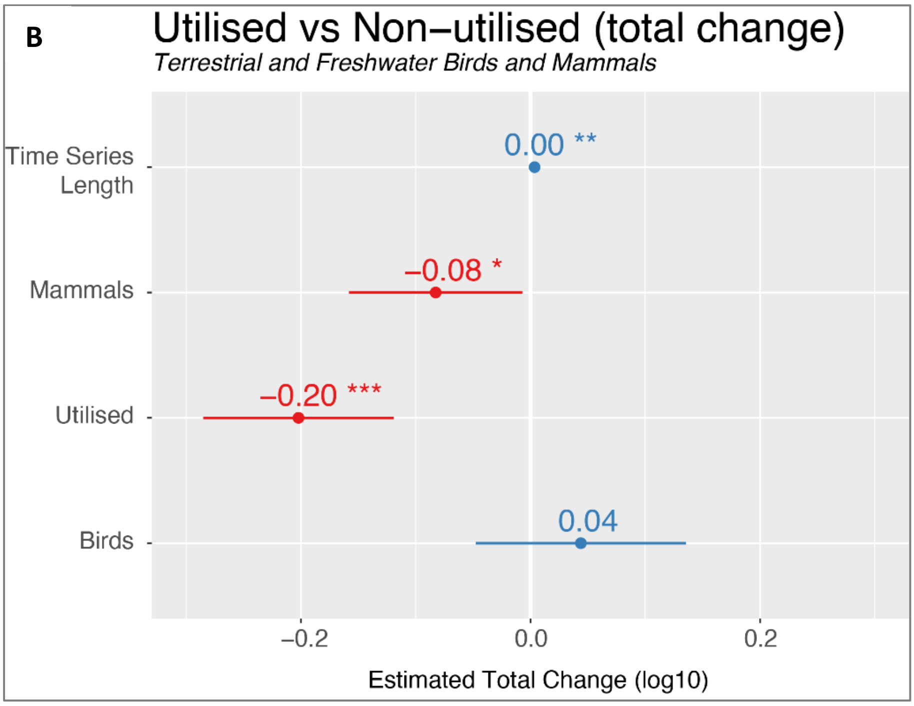


**Figure S6 Index of utilised and non-utilised populations for species of terrestrial and freshwater birds and mammals; Related to Figure 4. (A)** Here, between 1970 and 2016, on average, utilised populations had declined by 68% (0.20 - 0.51) and non-utilised populations had declined by 12% (0.71 - 1.09). **(B)** Estimated overall total change from the best linear mixed-effect model including Family, Binomial and location as random effects. Coefficients show the estimated overall change (log10) in each group. We found no significant interaction between taxonomic group and utilisation, with utilised populations of any taxa (*Utilised*) significantly more significantly more likely to be in decline.

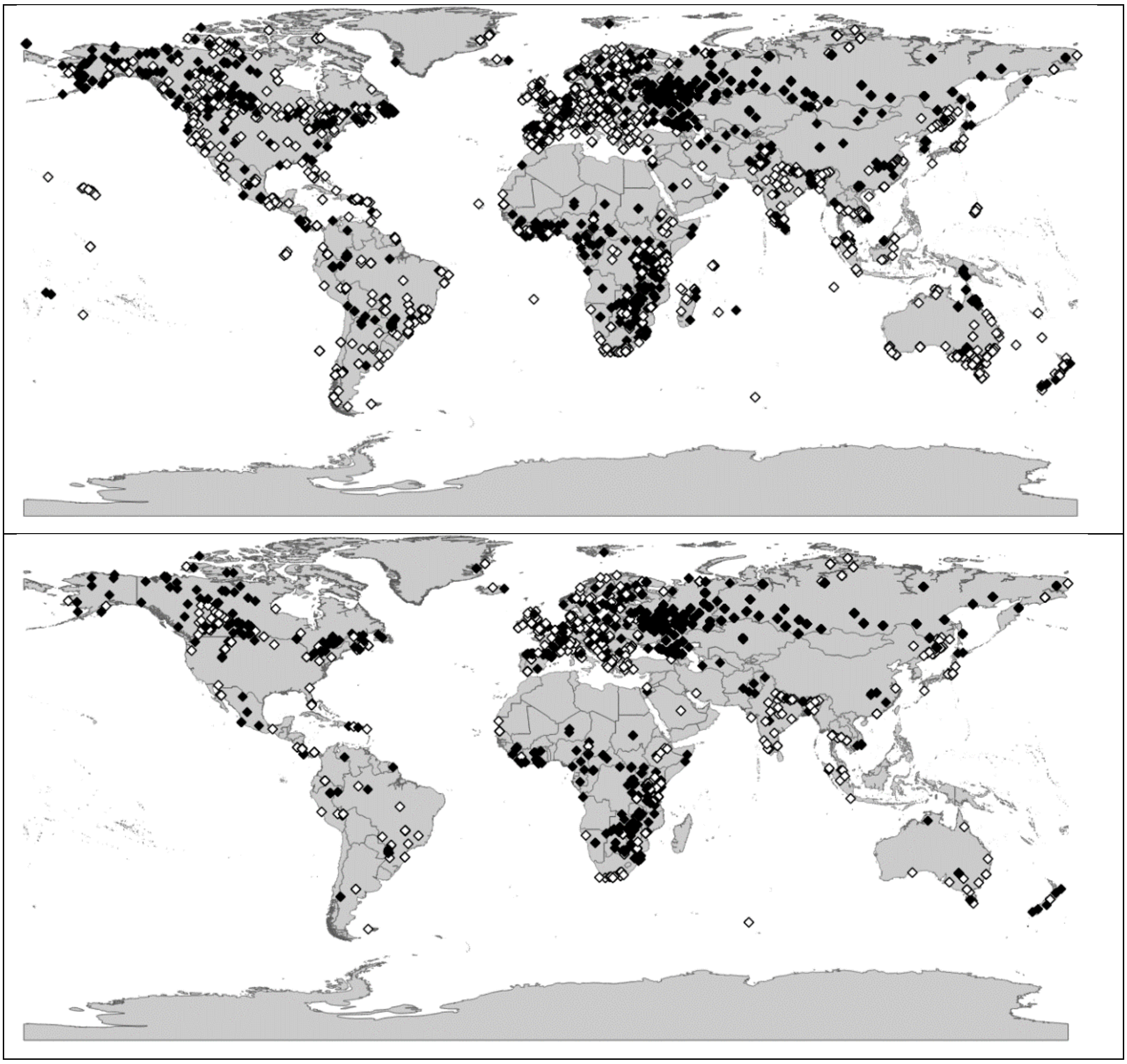

**Figure S7. Locations of utilised (black diamonds) and non-utilised (white diamonds) for terrestrial and freshwater birds, mammals and fish (A)** all species **(B)** the matched species of terrestrial and freshwater birds, mammals and fish

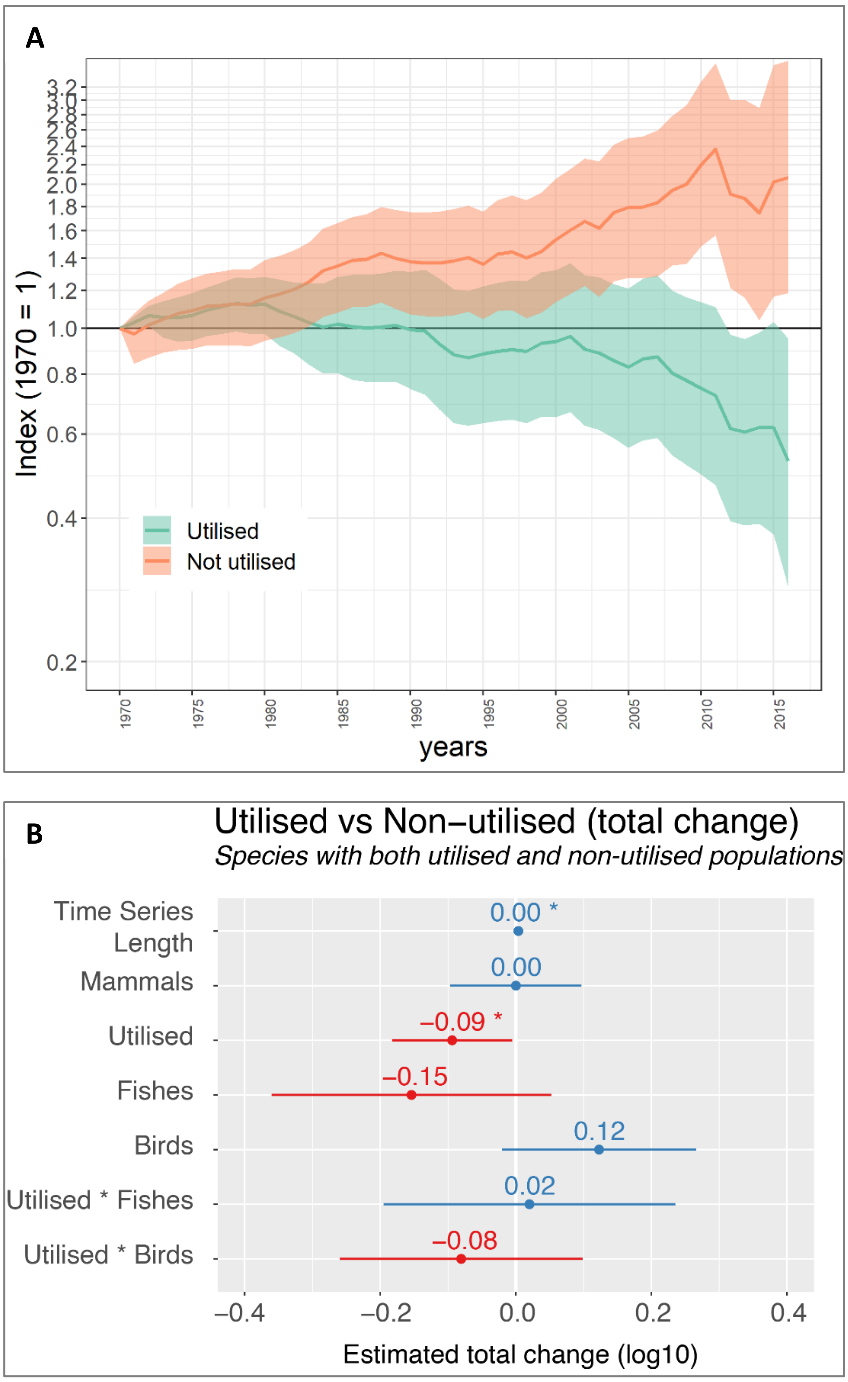


**Figure S8. Index of utilised and non-utilised populations for *matched* species of terrestrial and freshwater birds, mammals and fish; Related to Figure 4.** (i.e. species that have both utilised and non-utilised populations) **(A)** Here, between 1970 and 2016, on average, utilised populations had declined by 47% (0.29 - 0.95) and non-utilised populations had increased by 207% (1.18 - 3.63). **(B)** Estimated overall total change from the best linear mixed-effect model including Family, Binomial and location as random effects. Coefficients show the estimated overall change (log10) in each group. The impact of utilisation varied significantly with taxonomic group.

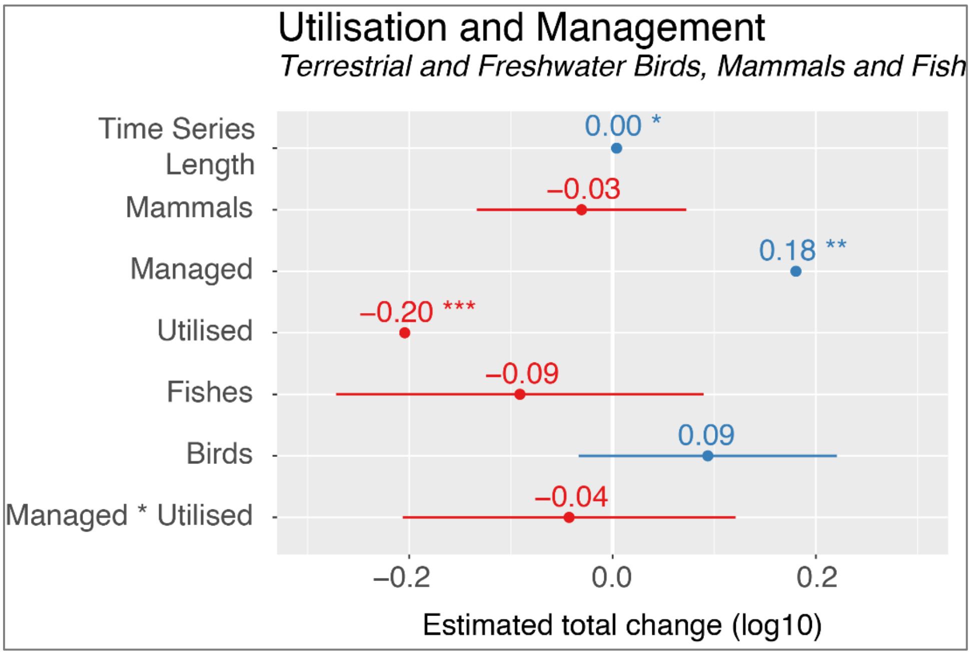


**Figure S9. Role of utilisation and management together.** For a limited number of species for which we had information on populations that were managed and unmanaged, we estimated overall total change from the best linear mixed-effect model including Family, Binomial and location as random effects. Coefficients show the estimated overall change (log10) in each group. The impact of utilisation varied significantly with management with utilised populations that were managed were significantly less likely to have declined.

**Table S6. Ancillary data fields used for disaggregation and modelling**

| Database field | Definition, coding and source of data | Examples or categories |
|---|---|---|
| Utilised | *Definition*: A population that is intentionally regularly or systematically utilised, either individuals or eggs. This may be sustainable or unsustainable, and the population does not necessarily have to be threatened by use or overexploited. This refers to consumptive use whereby individuals or parts of individuals are removed from the wild.<br>*Coding*: 'Yes', 'No', 'Unknown'<br>*Source*: This information was taken from the source of the population data | What is included:<br>Hunting (including subsistence, sport and trophy hunting)<br>Collecting<br>Fishing (consumptive including commercial, artisanal, sport, angling)<br><br>What is not included:<br>Wildlife tourism<br>Capture and release fishing<br>Education and research in situ<br>Viewing or experiencing for cultural or spiritual reasons |
| Managed | *Definition*: A population that receives targeted management (some of which involves sustainable use). This is usually to promote recovery in a population or can incentivise it's use for conservation. It can include measures to stem 'unsustainable' population growth.<br>*Coding*: 'Yes', 'No', 'Unknown'<br>*Source*: This information was taken from the source of the population data | What is included:<br>Supplementary feeding<br>Reintroduction<br>Captive breeding<br>Legal protection<br>Quotas for hunting<br>Provision of nest materials<br>Culling of predators of species being monitored<br>Culling of species being monitored (e.g. if overpopulated)<br><br>What is not included:<br>Protected area (unless it is specifically for that species – e.g. a tiger reserve) |
| Threats | *Definition*: A current threat that has been identified for the population, according to expert opinion.<br>*Coding*: Up to three threats coded per population<br>*Source*: This information was taken from the source of the population data | Climate change<br>Overexploitation (includes hunting and collecting, indirect killing, pet trade, sport hunting, persecuted as a pest)<br>Habitat loss/degradation<br>Invasive species and disease<br>Pollution |
| System | *Definition*: The system that best represents the location and habitat the population (not species) occupies. This is based largely on where the population was monitored and its primary habitat.<br>*Coding*: One system selected<br>*Source*: The location information is taken from the source of the | Terrestrial<br>Freshwater<br>Marine |

| | | |
|---|---|---|
| | population data. The primary habitat information is taken from Birdlife (birds), IUCN Red List (mammals, amphibians, reptiles), Fishbase (fish) | |
| IPBES region | *Definition*: Socio-political region defined by IPBES (IPBES 2015). If a population spans more than one region, the region containing the greater proportion of the location is selected.<br>*Coding*: One region selected<br>*Source*: (IPBES 2015) | Africa<br>Americas<br>Europe and Central Asia<br>Asia Pacific |
| IPBES subregion | *Definition:* Socio-political region defined by IPBES (IPBES 2015). If a population spans more than one subregion, the subregion containing the greater proportion of the location is selected.<br>*Coding:* One region selected<br>*Source:* (IPBES 2015) | Southern Africa<br>Central and Western Europe<br>North America |
| Geographical coordinates | *Definition*: The XY coordinates for the population – usually the centroid<br>*Coding*: Degrees, minutes and seconds or Decimal degrees<br>*Source*: This information was taken from the source of the population data or, if absent, an online geographical database | |
| Time-series length | *Definition*: The timeframe (number of years) from the first year the population was monitored to the final year. All intervening years were counted regardless of whether monitoring occurred in that year or not<br>*Coding*: Number of years<br>*Source*: This information was taken from the source of the population data | |

**Table S7. Indices of abundance calculated according to taxa, system and utilisation categories**

| | Utilised (All vertebrates) | Utilised and non-utilised (Birds, fish and Mammals) | Utilised and non-utilised (Birds and Mammals) |
|---|---|---|---|
| **Terrestrial and freshwater** | Global<br>IPBES regions<br>IPBES subregions | Global<br>Global – matched species only | |
| **Marine** | Global<br>IPBES regions<br>IPBES subregions | | |
| **All systems** | | Global<br>Global – matched species only | Global<br>Global – matched species only |
